# Characterization of Z-DNA dynamics across the tree of life

**DOI:** 10.64898/2025.12.30.697008

**Authors:** Georgios Megalovasilis, Eleftherios Bochalis, Michail Patsakis, Dionysios V. Chartoumpekis, Guliang Wang, Karen M. Vasquez, Ilias Georgakopoulos-Soares

## Abstract

Z-DNA/Z-RNA is an alternative left-handed nucleic acid conformation with established and emerging roles in gene regulation, immunity, and genome instability. However, its occurrence dynamics and lineage specificity across the tree of life have not yet been fully characterized. Utilizing the recently developed and improved Z-DNA searching tool, ZSeeker, we analyzed 281,139 complete organismal genomes, including multiple Telomere-to-Telomere genome assemblies, and generated genome-wide Z-nucleic acid maps, examined their topography, and compared them to dinucleotide-preserving controls. Cellular genomes featured pervasive Z-DNA enrichment relative to expectation, with enrichments of ∼1.5 and ∼1.7-fold in Bacteria and Archaea and ∼3-fold in Eukaryota. In contrast, Viruses exhibited large differences between lineages, with modest enrichment in several DNA viral groups and pronounced depletion across RNA clades, most notably Influenza A/B strains. We built a LASSO regression model trained on non-Influenza viruses (cross-validated R² ≈ 0.73), which identified GC content, genome type, and host type as the leading predictors for Z-nucleic acid density, yet it significantly over-predicted Z-RNA density in Influenza A/B. More than 99% of assemblies exceeded the +2 SD threshold, and a “typical Influenza” genome was predicted at 2.76 bp/kb compared to ∼0.016 bp/kb observed (a ∼170-fold overestimation based on chance alone). Together, these results reveal domain- and lineage-specific regimes: cellular genomes are enriched for Z-DNA consistent with regulatory roles, whereas influenza viruses appear to have undergone strong, lineage-specific depletion of Z-RNA-forming sequences, likely reflecting evolutionary pressure tied to host sensing pathways.

## Introduction

DNA primarily adopts the canonical right-handed B-form double helix, but it can also adopt alternative conformations collectively termed non-B DNA, each with distinct biological implications (Wang and Vasquez 2023). Z-DNA is a left-handed helical conformation characterized by a zigzag phosphate backbone and an alternating anti-syn arrangement of bases, typically forming in sequences with alternating purines and pyrimidines (Wang et al. 1979; Rich and Zhang 2003;Herbert A. 2019). The propensity to form Z-DNA varies among dinucleotides, with GC/CG repeats being the most favorable and AT/TA dinucleotides the least (Jovin et al. 1983; Peck and Wang 1983; Ho et al. 1986). Although the human genome is less GC-rich than prokaryotic genomes, Z-DNA-forming sequences are very abundant, with GT/AC repeats estimated to account for more than 0.25% of the human genome (Zhao et al. 2010). In addition to a favorable base sequence, Z-DNA formation is promoted by physiological factors such as negative supercoiling, high ionic strength, cytosine methylation, and is often transient in nature (Peck et al. 1982; Nordheim et al. 1982; Nordheim and Rich 1983; Azorin et al. 1983). Interestingly, it is energetically less costly than other non-B-DNA structures, and negative supercoiling during transcription alone is often sufficient to induce its formation (Shin et al. 2016; Subramani and Kim 2023; Lafer et al. 1981; Wang et al. 1979; Kouzine et al. 2017; Run et al. 2025; Schwartz et al. 1999). Although for many years Z-RNA, the RNA counterpart of Z-DNA, was not believed to form under natural conditions due to its less energetically favorable conformation, recent studies have identified Z-RNA duplexes as physiologically relevant structures (Balachandran and Mocarski 2021).

Detecting Z-DNA *in vivo* has historically been challenging due to its dynamic and transient nature, higher energy state, and thermodynamic instability, but the development of Z-DNA-specific antibodies and proteins containing high-affinity binding domains (e.g., Zα) has enabled *in vivo* detection, though each with certain limitations. *In vitro*, structural techniques such as circular dichroism, NMR, and X-ray crystallography have enabled its characterization (Shin et al. 2016; Subramani and Kim 2023; Lafer et al. 1981; Wang et al. 1979; Kouzine et al. 2017; Run et al. 2025; Schwartz et al. 1999). Based on experimental evidence, multiple computational methods have been developed to detect loci predisposed to Z-DNA formation (Peck and Wang 1983; Ho et al. 1986; Cer et al. 2013; Wang et al. 2025; Umerenkov et al. 2023). We recently developed ZSeeker, a computational tool for Z-DNA prediction that outperforms existing methods in accurately identifying Z-DNA-forming loci in genomic sequences (Wang et al. 2025), which can be leveraged to examine the topography of Z-DNA sites across genomes.

Research efforts have centered on elucidating the biological roles of Z-nucleic acid sequences, and Z-DNA formation has been associated with gene regulation, nucleosome positioning, and transcription initiation (Wong et al. 2007; Georgakopoulos-Soares et al. 2022; Schroth et al. 1992; Fang et al. 2024; Beknazarov et al. 2024). Loci predisposed to Z-DNA formation are preferentially distributed at gene silencers, ribosomal DNA arrays, and transcription start sites (Schroth et al. 1992; Wang et al. 2025). Functional roles of Z-nucleic acids in the modulation of innate immunity have also been uncovered, such as in inflammation and host cell death mediation during viral infections (Rothan et al. 2019). For adaptive immunity, a recent study showed that promoter-proximal Z-DNA loci serve as a marker, guiding autoimmune regulator (AIRE) to select genes for thymic T-cell tolerization by enhancing DNA double-strand breaks and promoter poising (Fang et al. 2024). Additionally, several proteins have been found to recognize Z-DNA and Z-RNA (Kim et al. 2000; Oh et al. 2002; Herbert A. 2019) via their specialized Zα domains, mediating many of their biological activities. For example, Zα domain-containing ADAR1 restrains Z-RNA-triggered ZBP1 (also a Z-binding protein) necroptosis, and releasing this block (e.g., with curaxin) sensitizes tumors to immune checkpoint therapy (Zhang et al. 2022). During viral infections (such as Influenza A and SARS-CoV-2), viral replication gives rise to Z-RNA structures that are sensed by the host protein ZBP1, which in turn activates RIPK3-dependent necroptosis to eliminate infected cells and stall viral spread (Zhang et al. 2020; Balachandran and Mocarski 2021). Together, these studies indicate that Z-DNA/Z-RNA in microbes shape genome stability and host-pathogen interactions, warranting a broader systematic study.

Loci predisposed to Z-DNA formation are associated with genomic instability (Bacolla et al. 2016; Georgakopoulos-Soares et al. 2018; Wang et al. 2006) and aging (Li et al. 2025), while the transcription and stability of several disease-related genes, including oncogenes, are also regulated by Z-DNA in humans (Wang et al. 2006; Georgakopoulos-Soares et al. 2018; McKinney et al. 2020; Fang et al. 2024). Z-DNA structures can render surrounding DNA regions prone to damage and cleavage, causing local genetic instability such as deletions and rearrangements (Wang and Vasquez 2007). This rapid evolution driven by double-strand breaks at Z-DNA loci underlies the repeated loss of pelvic structures in stickleback fish (Xie et al. 2019), indicating roles associated with adaptation. These findings suggest that Z-DNA links structural plasticity to both pathogenic mutagenesis and evolutionary change.

Examination of Z-DNA in Prokaryota and Viruses remains limited, with few studies to date (Freund et al. 1989; Wang et al. 2023; Kim et al. 2003; Buzzo et al. 2021). It has been shown that Z-DNA formation in bacteria results in genomic instability (Freund et al. 1989; Wang et al. 2023; Kha et al. 2010) and that extracellular Z-DNA formation assists in the development of bacterial biofilms (Buzzo et al. 2021). The recent rapid increase in available organismal genome assemblies spanning the tree of life offers an unprecedented opportunity to explore genome architecture. Yet, the distribution and functional significance of Z-DNA across diverse organisms remains poorly understood. Here, we conducted a comprehensive analysis of Z-nucleic acid-forming sequences across 281,139 genome assemblies spanning the tree of life. We report large differences in Z-nucleic acid density and enrichment among the 3 domains of life and Viruses, as well as in the distribution of Z-DNA in different genomic sub-compartments. Strikingly, we also identify a specific viral group (Orthornavirae kingdom; Influenza A/B) to be completely depleted of Z-RNA sequences, and overestimated for Z-RNA density by our model trained on non-Influenza viruses.

## Results

### Mapping of potential Z-nucleic acid-forming sequences in organismal genomes

We examined 281,139 organismal genomes (representing 106,125 unique species) across the tree of life for the prevalence of Z-DNA-forming and Z-RNA-forming sequences, including 209,871 viral assemblies (featuring 86,396 unique species), 69,612 bacterial (18,551 species), 827 archaeal (669 species), and 829 eukaryotic assemblies (509 species). Z-nucleic acid sequences were predicted by the ZSeeker software (Wang et al. 2025), using the nucleotide sequence of each complete genome as input. In total, we identified 854,794,975 potential Z-nucleic acid-forming sequences across all organismal genomes. The average Z-nucleic acid-forming sequence density was calculated at 53.7 bps per kilobase (kb). The density of motifs, however, varied substantially among species, ranging from 0 to 778 bps per kb. An uncultured beta proteobacterium strain (belonging to the Betaproteobacteria class) featured the highest Z-DNA density (∼778 bps/kb), with an ∼14.5-fold enrichment compared to the average density of all genomes, followed by different strains of *Cellulomonas iranensis* (∼774.5 bps/kb). Regarding ZSeeker metrics, the longest Z-DNA sequence was found to be 65,589 bps in *Neophocaena sunameri*, and the highest-scoring one was found in *Electrophorus electricus*. Surprisingly, in 154,629 viral, 194 bacterial, and 12 eukaryotic genomes, we found no predicted Z-DNA sequences at all. Highest-ranking species for Z-nucleic acid-forming sequence density, lengths, and scores can be found in the **Suppl. Tables 1 & 2**.

We report significant differences in Z-nucleic acid-forming sequence density among the three domains of life and Viruses (also referred to as the four superkingdoms), with Bacteria exhibiting the highest (161.4 bps per kb), followed by Archaea (106.7 bps per kb), Eukaryota (39.2 bps per kb), and Viruses (17.9 bps per kb) (**Fig. 1a**). Next, we examined differences in Z-nucleic acid-forming sequence density with finer taxonomic resolution, including kingdoms and phyla. We observe that at the kingdom level (**Fig. 1b**), Pseudomonadati and Helvetiavirae displayed the highest densities, with raw values equal to 189.2 and 179.4 bps per kb, respectively. Kingdoms ordered by raw mean values are featured in **Supp. Fig. 1a**. Conversely, the lowest mean densities are observed in Orthornavirae and Fusobacteriati, with 1.2 and 0.2 bps per kb. At the phylum level (**Fig. 1c**), we observe even greater variation. The bacterial phylum Vulcanimicrobiota shows the highest Z-DNA density (504.4 bps per kb), whereas the viral phylum Negarnaviricota has the lowest (33.1 bps per kb). These findings indicate striking domain- and lineage-specific heterogeneity in the frequency of Z-DNA sequences across the tree of life.

**Figure 1:**
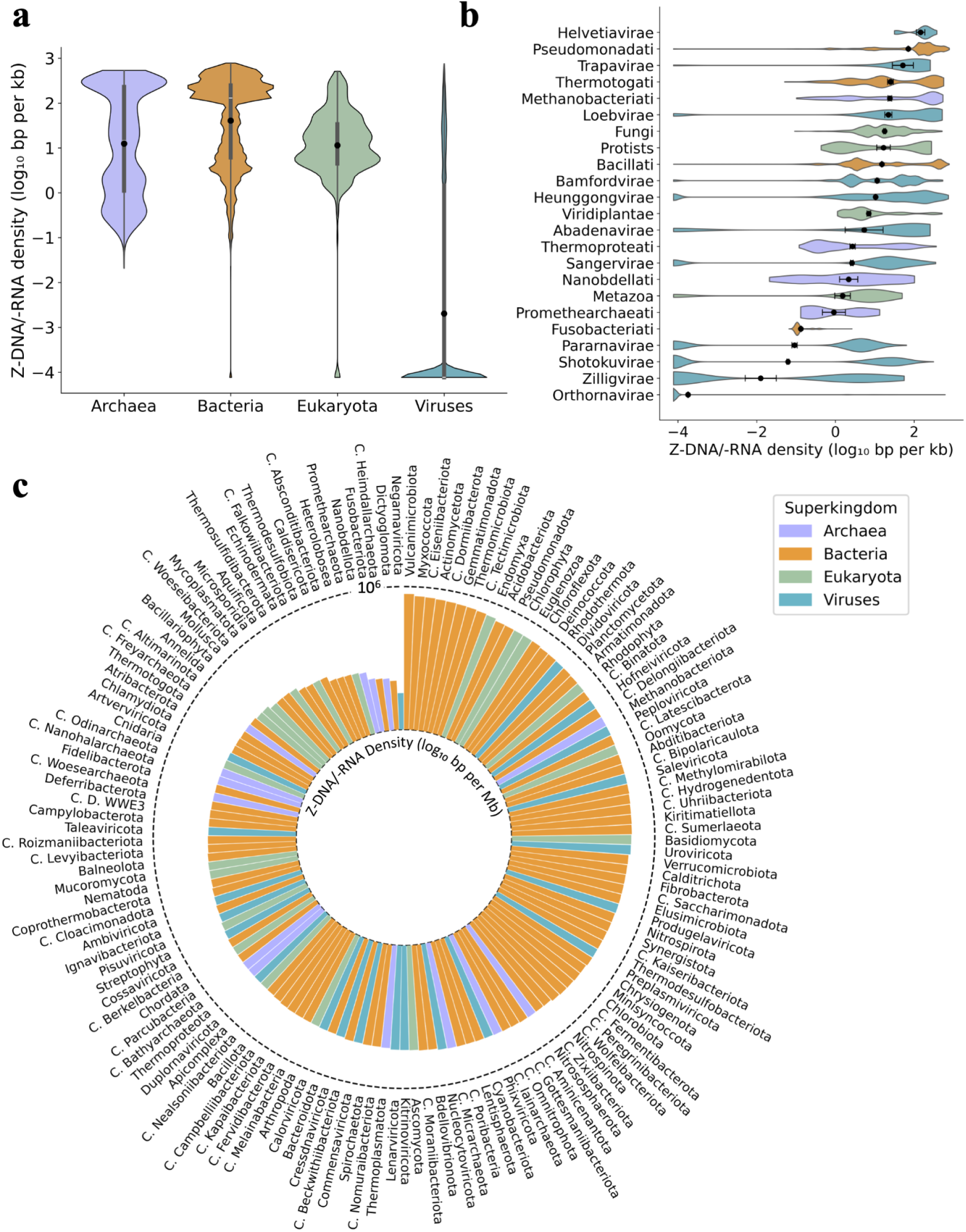
Average Z-nucleic acid sequence density across taxonomic levels. Average Z-DNA density (in bps per kb) across **a.** the four superkingdoms, and **b.** kingdoms, visualized with violin plots. **c.** Average Z-DNA/Z-RNA density (in bps per Mb) across phyla. Black dots indicate group means, with error bars representing the standard error of the mean (SEM). Colors denote superkingdom classification.

### Genome composition and size influence Z-nucleic acid-forming motif density

To explore further differences in Z-nucleic acid-forming motif dynamics among the four superkingdoms, we compared the distributions of four metrics: the mean and maximum Z-nucleic acid-forming sequence prediction score and the mean and maximum predicted length (**Fig. 2a**). The Z-nucleic acid-forming sequence score indicates the likelihood that a sequence may adopt a Z-nucleic acid conformation. Viruses consistently displayed the lowest values across all four metrics, with extremely large effect sizes (r > 0.8 for mean and r > 0.9 for maximum metrics). This was confirmed with Mann-Whitney U tests with Bonferroni correction (adjusted p-value < 0.0001). Among the domains of life, Bacteria and Eukaryota showed no difference in mean score (adjusted p-value = 0.35), but Bacteria had slightly shorter mean predicted Z-DNA segments (r = 0.31) and modestly lower maximum values (r ≤ 0.15). Archaea differed from both Bacteria and Eukaryota across most metrics (adjusted p-value < 0.0001), but with more modest effect sizes (|r| ∼ 0.1 - 0.5): Archaea tended to have lower mean scores and lengths than Bacteria, but higher maximum values compared to both Bacteria and Eukaryota. Given the sample sizes, these comparisons, while statistically significant, may indicate only small quantitative shifts among the domains of life. Motivated by these differences, we next sought to identify the determinants, including genome architecture and composition, and the extent to which they can explain the observed disparities in Z-nucleic acid sequence density across lineages.

**Figure 2:**
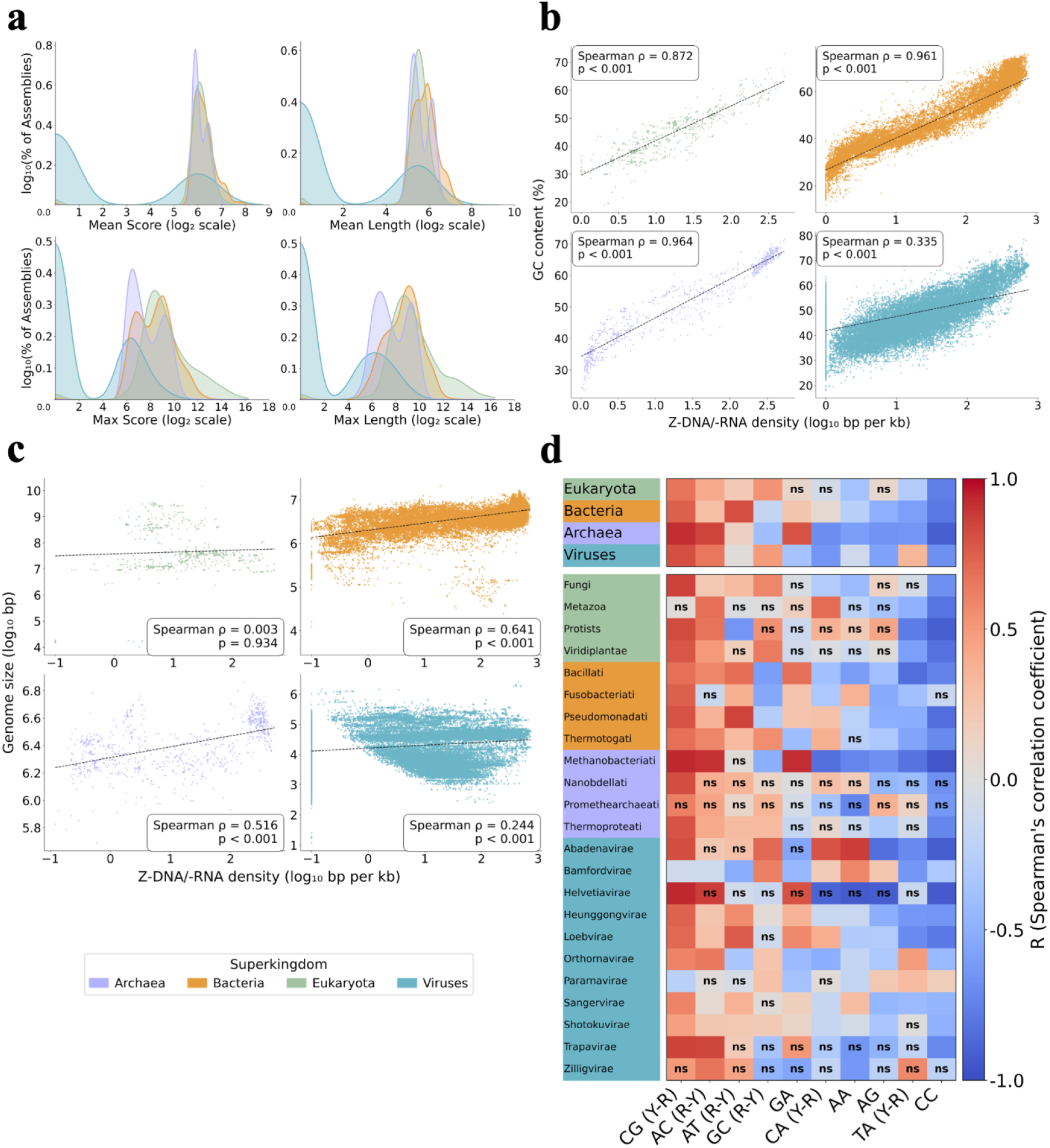
Comparative landscape of Z-DNA/-RNA sequences and genomic correlates across superkingdoms. **a.** Mean and max Z-DNA/RNA score and length distributions of superkingdoms. Correlation between **b.** GC content, **c.** genome size, **d.** dinucleotide frequencies and Z-DNA/RNA density across superkingdoms.

Therefore, we investigated whether there is a correlation between GC content and Z-nucleic acid sequence density (**Fig. 2b**) and between genome size (**Fig. 2c**) in any of the four superkingdoms. For GC content, we find that Archaea, Bacteria, and Eukaryota display very high positive correlations (Spearman correlations, ρ ∼ 0.97, ρ ∼ 0.96, and ρ = 0.87, respectively), whereas Viruses show a positive but much weaker correlation (Spearman correlation of ρ ∼ 0.34) (all p < 0.001). This pattern, however, is not unexpected, as Z-nucleic acid-forming sequences are known to most favorably form in regions with alternating purine-pyrimidine motifs, especially GC-rich repeats (Krall et al. 2023; Sahayasheela et al. 2025). We also compared genome size and Z-nucleic acid-forming sequence density to evaluate whether larger genomes tend to feature higher densities. All superkingdoms exhibit a positive correlation between genome size and Z-nucleic acid-forming sequence density, with stronger correlations in Bacteria (ρ = 0.66) and Archaea (ρ = 0.54), and weaker correlations in Eukaryota (ρ = 0.26) and Viruses (ρ = 0.19) (all p < 0.001).

Because GC content strongly influences dinucleotide frequency composition in organismal genomes (Forni et al. 2024), we accounted for the effect of overall base composition from specific sequence preferences. To do this, we calculated enrichment scores by comparing observed dinucleotide frequencies to the expected values given the mononucleotide composition of each genome. This approach controls for GC content and allows us to detect whether certain dinucleotides are over- or under-represented beyond what would be predicted from base composition alone. Based on this notion, we examined whether variation in dinucleotide enrichment helps explain differences in Z-nucleic acid-forming sequence density. For each superkingdom and kingdom, we computed Spearman correlations between dinucleotide enrichment (observed/expected ratio, controlling for G+C content) and log-transformed Z-nucleic acid-forming sequence density. We observe that almost half of all dinucleotides were non-significant, especially for Archaea and Eukaryota that feature the fewest assemblies. A clear pattern, however, emerges among the remaining cases: Y-R (or R-Y) dinucleotides (pyrimidine-purine pairs such as CG, AC) mostly exhibited positive correlations with Z-nucleic acid-forming sequence density across multiple taxon levels. In contrast, non-Y-R dinucleotides were less often positively correlated, and when significant, tended to exhibit negative correlations (**Fig. 2d**). This indicates that, beyond the influence of GC content, local enrichment of alternating Y-R motifs acts as a driver of Z-nucleic acid-forming sequence formation potential.

### Benchmarking Z-nucleic acid sequence density patterns against matched simulations

To better assess Z-nucleic acid sequence enrichment, for every genome, we created a simulated matched control genome that preserved the underlying dinucleotide composition and length. Across the 281,139 genomes analyzed, 114,033 had at least one Z-nucleic acid prediction in both observed and simulated genomes, while 12,271 had predictions only in the observed genomes, indicative of potential enrichment. Conversely, 28,003 genomes had predictions only in their simulated counterparts, suggesting depletion. The remaining 126,832 genomes showed no predicted Z-nucleic acid sequences in either their observed or simulated genome (**Fig. 3a**). Notably, genomes without any Z-nucleic acid predictions, as well as those showing predictions only in the observed or only in the simulated versions, comprised mostly viral assemblies.

**Figure 3.**
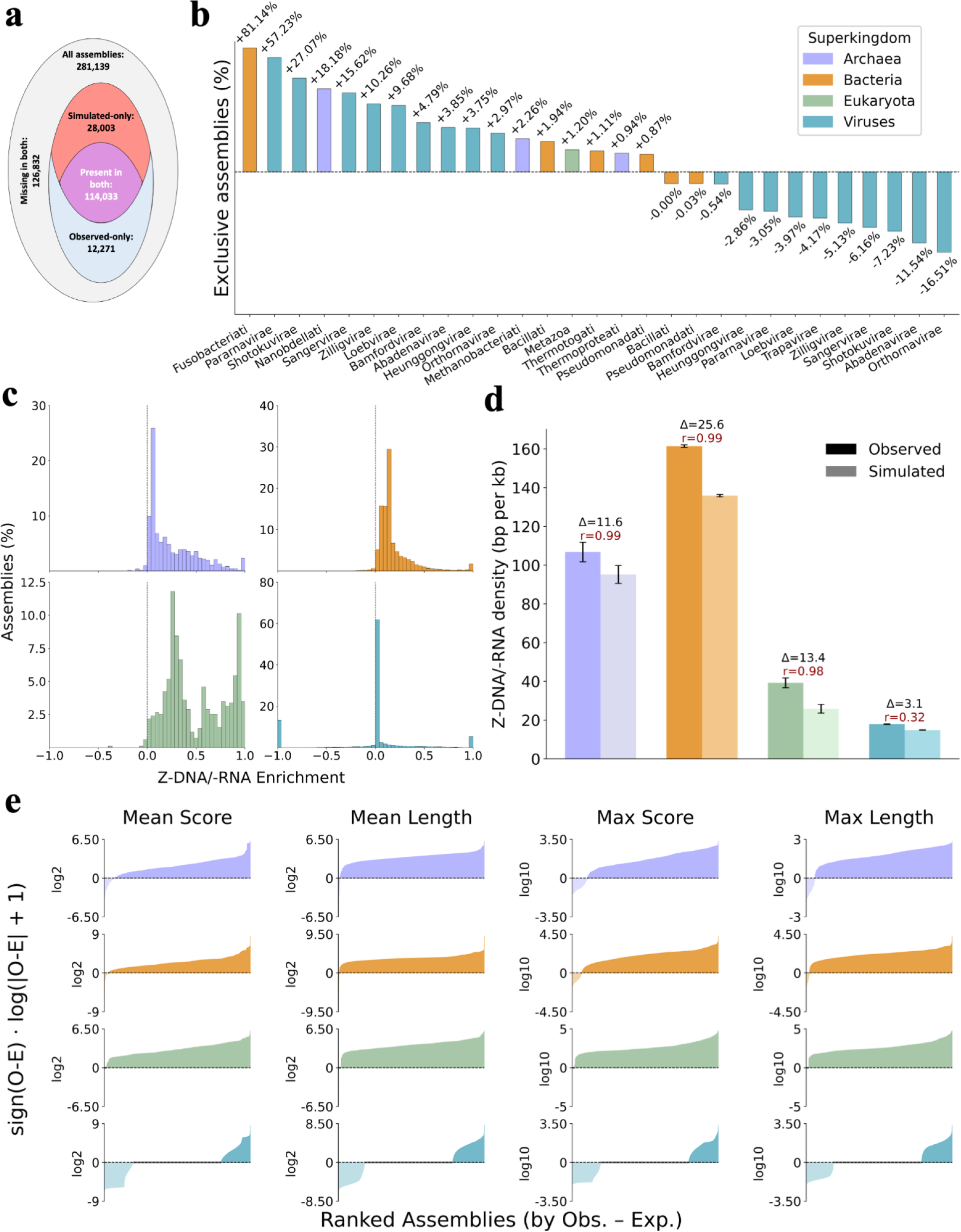
Observed versus simulated Z-nucleic acid sequence densities across genomes. **a.** Numbers of genomes with no predicted Z-nucleic acid tracts or at least one predicted tract in observed and shuffled controls. **b.** Percentages of assemblies within each kingdom showing exclusive presence of Z-nucleic acid predictions in observed or shuffled genomes. **c.** Normalized enrichment scores of Z-DNA/Z-RNA densities within each superkingdom (−1 = complete depletion, +1 = maximal enrichment). **d.** Z-nucleic acid density pairwise comparisons between observed and control genomes by superkingdom. **e.** Z-nucleic acid tracts’ length and score enrichment across superkingdoms.

When examining exclusive assembly presence (referring to none versus at least one Z-nucleic acid prediction) in original versus simulated genomes by kingdom, differences were mostly driven by viral groups (**Fig. 3b**). Percentages reported below refer to the proportion of assemblies within each kingdom. Several viral kingdoms showed strong enrichment of observed-only assemblies, including Pararnavirae (+57.2% versus -3.1%), Shotokuvirae (+27.1% versus -7.2%), and Sangervirae (+15.6% versus -6.2%). In contrast, Orthornavirae exhibited a striking depletion, with only +3.0% observed-only compared to -16.5% shuffled-only control assemblies, representing a net deficit of 21,053 assemblies. Among organismal assemblies, modest enrichment was observed in groups such as Bacillati (+1.9%) and Metazoa (+1.2%). Smaller kingdoms such as Abadenavirae (+3.9% versus -11.5%) and Trapavirae (-4.2%) also showed depletion, although these patterns should be interpreted cautiously given the limited number of genomes available. Similarly, extremely high percentages in kingdoms with very low Z-nucleic acid sequence density initially (e.g., Fusobacteriati, +81.1%) reflect division-by-zero effects, stemming from enrichment calculation, rather than clear kingdom trends.

To directly compare enrichment and depletion of Z-DNA/Z-RNA-forming sequences among superkingdoms, we calculated these values, considering observed and shuffled (control) genomes (**Fig. 3c**). We find that Archaea, Bacteria, and Eukaryota consistently exhibit higher levels of Z-DNA-forming sequences than expected for almost all of their assemblies. On average, Eukaryota show a ∼3-fold enrichment, Archaea a ∼1.7-fold enrichment, and Βacteria a ∼1.5-fold enrichment of Z-DNA-forming sequences relative to shuffled controls, with >97% of assemblies enriched in each group. In contrast, viral assemblies featured only ∼19% enriched and ∼18% depleted assemblies. However, the average enrichment score was slightly negative, indicating that when depletion occurred it tended to be stronger in magnitude than the enrichment cases. Pairwise Mann–Whitney U tests confirm that Eukaryota are significantly more enriched than Archaea and Bacteria, and that all three cellular superkingdoms differ markedly from Viruses (Bonferroni-corrected p < 0.005). To explore these patterns at finer taxonomic resolution, we examined enrichment across individual kingdoms. **Supp. Fig. 1b** displays enrichment graphs for kingdoms with a number of assemblies >80. Fusobacteriati displayed the strongest enrichment, with a striking ∼30-fold change, compared to ∼1.3-1.6-fold of the other bacterial kingdoms. Archaeal kingdoms feature a notable enrichment of around 1.5-3-fold, while all eukaryotic kingdoms are highly enriched: Viridiplantae (∼9-fold), Metazoa (∼6-fold), Protists (∼3.2-fold), and Fungi (∼2.2-fold). Among viral kingdoms, most showed a low to moderate enrichment, with Bamfordvirae (∼2.8-fold) and Pararnavirae (∼3.5-fold) exhibiting the highest. In contrast, the viral Orthornavirae kingdom was globally the only one depleted (∼0.75-fold), while it comprises >150,000 assemblies.

To investigate enrichment differences within each superkingdom, we performed paired Wilcoxon signed-rank tests (due to paired data and a non-normality assumption) and estimated rank-biserial correlation effect sizes **(Fig. 3d)**. The results confirmed highly significant enrichment of Z-DNA across Archaea (r = 0.98), Bacteria (r = 0.99), and Eukaryota (r = 0.98), with large effect sizes (all q < 0.001, where q values are FDR-adjusted p-values). In contrast, although the viral dataset also yielded a formally significant result (q < 0.001), the effect size was small (r = 0.32) and the mean difference in Z-nucleic acid sequence density between observed and shuffled genomes was only ∼3 bps (compared to 12-26 bps for the domains of life), suggesting little biological relevance and consistent with the overall lack of enrichment in Viruses.

### Enrichment of Z-nucleic acid tract length and score relative to shuffled controls

We next characterized mean and maximum values of Z-nucleic acid tract lengths and ZSeeker scores between genomes and their matching simulated control genomes (**Fig. 3e**). Within each row (superkingdom), assemblies are ranked left-to-right by the observed - expected difference for the metric. Values above the baseline indicate the observed genome exceeds its control (enrichment), while the ones below indicate that the control exceeds the observed (depletion). We log2-transformed the mean and log10 the max metrics to compensate for their different ranges and enhance comparability. The paired Wilcoxon signed-rank tests with rank-biserial effect sizes and FDR-corrected q-values agree with the visual pattern. For mean scores, effects are large in Archaea (r = 0.86, q-value < 0.001), Bacteria (r= 0.95, q-value < 0.001), and Eukaryota (r = 0.99, q-value < 0.001), whereas Viruses show a small negative effect (r = −0.09, q-value < 0.001, expected > observed). For mean length, Archaea (r = 0.98, q-value < 0.001), Bacteria (r = 0.99, q-value < 0.001), and Eukaryota (r = 0.99, q-value < 0.001) are once again strongly enriched, while Viruses are none to slightly negative (r = −0.04, q-value < 0.001). Their maximum counterparts, however, behave somewhat differently: max length is strongly positive in all four superkingdoms (r = 1.00, q-value < 0.001), and max score shows large effects in cellular groups: Archaea (r = 0.80, q-value < 0.001), Bacteria (r = 0.87, q-value < 0.001), Eukaryota (r = 0.99, q-value < 0.001), with only a very small effect in Viruses (r = -0.04, q-value < 0.001).

### Z-nucleic acid landscape in genomic compartments and transcription start and end sites

To investigate putative regulatory roles of Z-nucleic acid-forming sequences, we next analyzed their enrichment in genomic sub-compartments across taxonomic groups. We examined differences in the enrichment of Z-nucleic acid-forming sequences, focusing on genes, coding sequences (CDS), and exons. Only assemblies with at least 1 potential Z-nucleic acid-forming sequence were included, as for genomes with 0 bps per kb Z-DNA/-RNA density there were no predictions to be mapped. **Figure 4a** displays enrichment scores for superkingdoms and phyla and for visualization purposes, it shows only phyla with an enrichment greater than or equal to 4X or lower than or equal to 0.25X (depletion). No viral phyla passed this threshold and therefore are not present in the panel. At the superkingdom level, we report very subtle differences, with Bacteria showing slight enrichment for all 3 features and Eukaryota exhibiting a depletion of Z-DNA sequences of 0.36 in exons. Examining differences in the phylum level reveals highly variant enrichment scores among taxa and genomic features. For genes, 2 eukaryotic phyla feature the highest (4.82X for Chordata, followed by some bacterial phyla) and lowest (0.23X for Annelida) enrichment. For CDS, Candidatus Altimarinota displays the highest score (43.75X), followed by Chordata (26X) and Fusobacteriota (5.62X). The remaining eukaryotic phyla show the most depletion (∼0.2X), followed by the archaeal Candidatus Heimdallarchaeota (0.36X). For exons, we report by far the most outliers, with the majority of phyla comprising very high or very low enrichment scores. Dictyoglomota (∼49X), Candidatus Altimarinota (∼44X) and Chordata (∼24X) constitute the highest-enrichment outliers, while the bacterial Kiritimatiellota, Chrysiogenota and Myxococcota (≤0.1X) show the greatest depletion.

**Figure 4:**
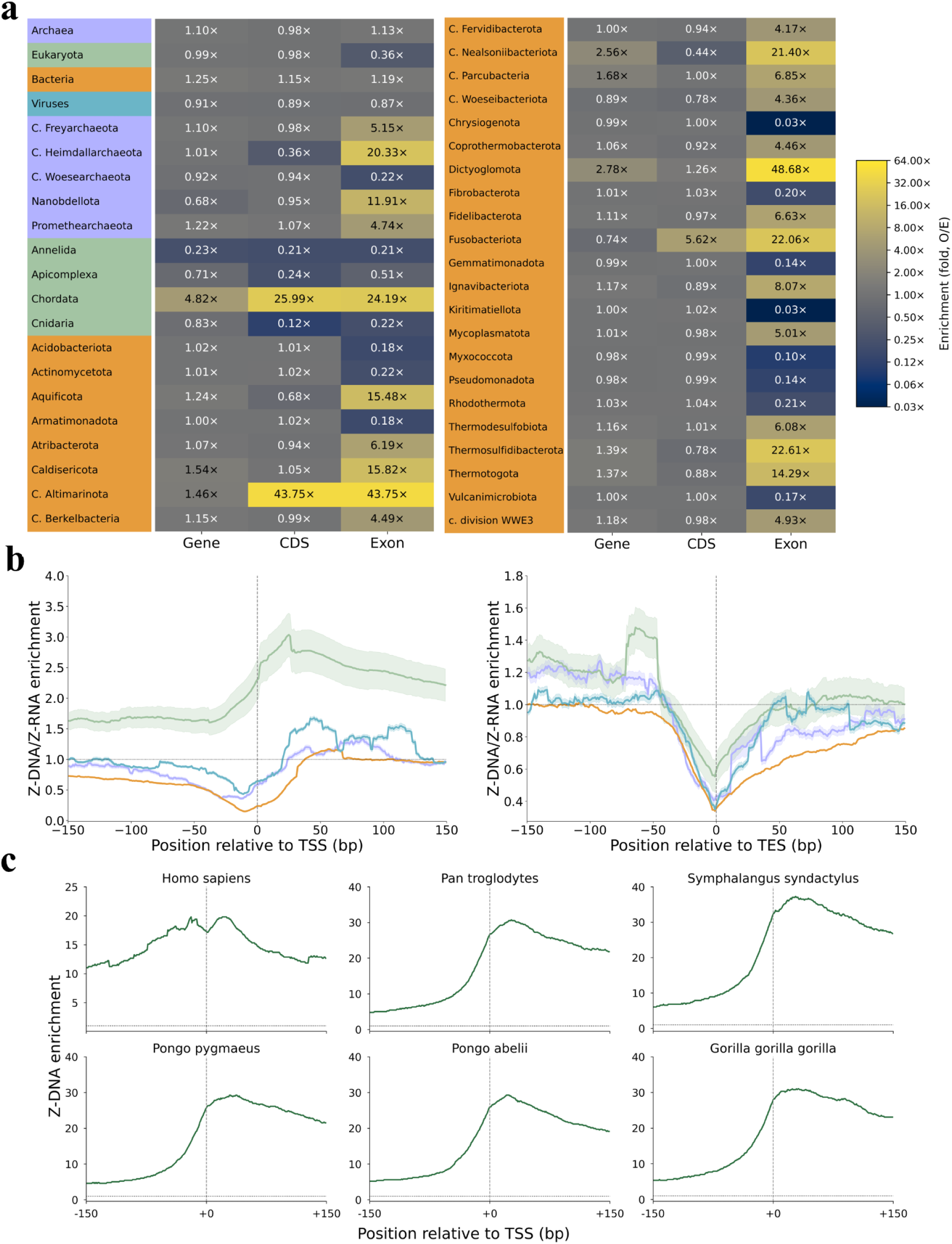
Cross-taxa Z-nucleic acid sequence analysis of genomic features and transcription start/end sites. **a.** Log₂ fold-enrichment of Z-nucleic acid sequences within genomic subcompartments in superkingdoms and phyla. **b.** Log₂ fold-enrichment of Z-nucleic acid sequences around TSSs and TESs by superkingdom. **c.** Log₂ fold-enrichment of Z-DNA around TSS in 6 T2T primate species and the human T2T assembly.

Since Z-nucleic acid sequence predictions were unevenly distributed across these genomic subcompartments, we next examined their distribution in relation to transcription start sites (TSSs) and transcription end sites (TESs), among the 4 superkingdoms (**Fig. 4b**). Pairwise Wilcoxon signed-rank tests with Benjamini-Hochberg FDR correction reveal that Eukaryota is the most enriched group, followed by Viruses. The lowest enrichment is observed in Bacteria and Archaea, with Archaea slightly above Bacteria (Wilcoxon, FDR-corrected q-value < 0.001 and large effect sizes r ≈ 0.8 for all pairs). TES comparisons yield similar results, besides Archaea and Viruses displaying a weaker statistical difference (small effect size r ≈ 0.16) and marginally significant (q-value ≈ 0.0057). Next, we utilized the genomes from the Telomere-to-Telomere (T2T) Primates consortium (Yoo et al. 2025) genomes, and compared the Z-DNA sequence enrichment around TSSs in the *Homo sapiens* assembly and 6 non-human primate genomes including *Gorilla gorilla* (gorilla), *Pan troglodytes* (chimpanzee), *Pongo abelii* (Sumatran orangutan), *Pongo pygmaeus* (Bornean orangutan) *Symphalangus syndactylus* (Siamang gibbon) (**Fig. 4c**) and *Pan paniscus* (bonobo) (**Supp. Fig. 1c**). Z-DNA enrichment around TSSs differed significantly across primates (Wilcoxon test; FDR q < 0.001 for all but one comparison; r ≈ 0.36–0.87). *Symphalangus syndactylus* showed the strongest enrichment, exceeding all other species, followed by *Gorilla gorilla gorilla*. *Homo sapiens* exhibited significantly lower overall TSS-proximal enrichment than all non-human primates, including *Pan paniscus*, but uniquely displayed elevated upstream enrichment (>10-fold), whereas other primates showed a sharp increase centered at the TSS. Among great apes, *Pan troglodytes* and *Pan paniscus* differed strongly, while *Pongo abelii* and *Pongo pygmaeus* were most similar; the latter comparison was the weakest, and *P. abelii* did not differ significantly from *P. paniscus*.

### Sparse regression identifies influenza viruses as drivers of Z-RNA absence

Given the consistent depletion patterns observed in some viral genomes, we next asked which viral lineages are depleted for Z-nucleic acid sequences and which genomic or ecological attributes best predict this depletion. We examined the Z-DNA/Z-RNA density across different viral groups, taking into account their genome and host types. Out of our total 209,871 viral assemblies, we extracted NCBI metadata for 148,098, with the rest missing well-curated information about either genome type (DNA or RNA), genome strand structure (single- or double-stranded) or host type (Eukaryota- or Prokaryota-infecting viruses). Dividing viral genomes into groups based on these features, we observe that bacterial ssDNA viruses showed the highest Z-nucleic acid sequence density (146.6 bps per kb), followed by ssDNA Archaea-infecting viruses (143.8 bps per kb) and bacterial dsRNA viruses (135 bps per kb) (**Supp. Fig 1d**). We also find that eukaryotic ssRNA viruses (the most abundant group, n = 140,191) exhibit the lowest density (0.19 bps per kb), as the majority of them (98.75% of genomes) display a complete Z-nucleic acid sequence depletion, lacking even a single prediction. When examining further which viral species this group comprised, we find they mostly include the RNA Influenza A and B viruses of the Negarnaviricota phylum (**Fig. 5a**). Pairwise comparisons between groups confirmed the highest differences in Z-nucleic acid sequence densities occur between the influenza viruses and all other viral types (Mann–Whitney U tests with Bonferroni corrections, p < 0.001 for all pairwise comparisons, with most effect size metrics ranging from 0.25 to 0.4). When calculating Z-RNA sequence enrichment relative to simulated control genomes within the Riboviria realm, only 0.27% of influenza genomes show enrichment, in contrast to 41.7% among the non-influenza Riboviria viruses, with 17.7% of influenza genomes being depleted instead (**Fig. 5b**).

**Figure 5.**
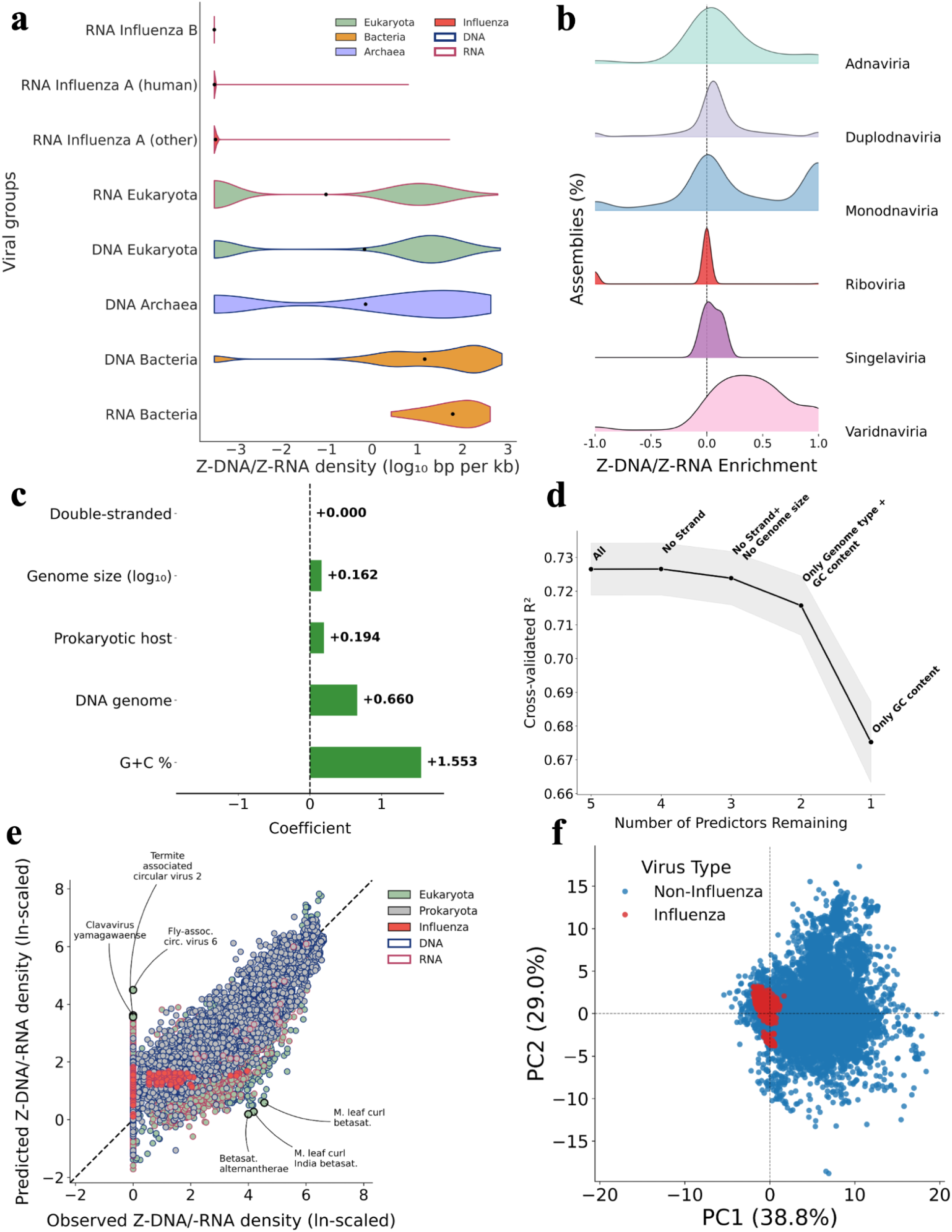
Distribution of Z-nucleic acid-forming sequences across viral genomes. **a.** Violin plot of Z-DNA/Z-RNA densities across major viral groups. **b.** Z-DNA/Z-RNA enrichment distributions by viral realm. **c.** LASSO regression coefficient analysis of non-influenza viruses. **d.** Strength of model predictors, as summarized by iterative removal of the weakest. **e.** LASSO model across all Viruses: over- and under-predictions **f.** Principal component analysis in the dinucleotide enrichment space visualizes Influenza clusters.

Next, we aimed to understand whether this massive Z-RNA-predicted sequence depletion in Influenza viruses can be attributed to common features in this group (taxonomy, short range of genome size and G+C content, ssRNA genome, eukaryotic hosts) or reflects an influenza-specific evolutionary pressure. We used LASSO (Least Absolute Shrinkage and Selection Operator) regression to generate a predictive model and assess how much these viral characteristics account for the observed variation in Z-DNA/Z-RNA densities (**Fig. 5c**). As the training set we defined the entirety of curated non-influenza viruses and as predictors we evaluated genome size, GC content, type of nucleic acid and strand structure, and host type. The shrinkage parameter (λ) was optimized using 5-fold cross-validation, selecting the value with the lowest prediction error. The cross-validated R² averaged 0.73 ± 0.01, showing that the model accounted for ∼73% of the variance in Z-DNA/Z-RNA densities across non-influenza viruses. The LASSO model identified GC content, DNA genome, and prokaryotic hosts as the strongest positive predictors of Z-DNA/Z-RNA density, with smaller effects for genome size and no effect for strand structure. Positive coefficients correspond to higher predicted densities. Iterative removal of the predictor with the lowest absolute coefficient showed that model performance remained stable (cross-validated R² = 0.72 - 0.73) until the 2 variables with the highest influence remained (GC content and genome type). After genome type was removed, performance declined to ∼0.675 (**Fig. 5d**).

After training on non-influenza viruses, the LASSO model was used to predict the Z-DNA/Z-RNA density each virus would be expected to have given its genome size, GC content, genome type, strand structure, and host. Systematic over- or under-estimation thus indicates densities that cannot be explained by these features alone. The regression model was applied to all curated viral genomes and its predictions for Z-DNA/RNA sequence densities were compared to the observed values (**Fig. 5e**). Influenza A (both human and non-human) genomes’ densities were overwhelmingly overestimated by the model, with more than 99% of assemblies falling above the +2 SD outlier threshold. Among non-influenza viruses, DNA Eukaryota-infecting ones were also strongly overestimated, with 29.1% of their assemblies falling beyond +2 SD, followed by RNA Eukaryota (non-influenza) (21.6%) and DNA Prokaryota (10.2%) viruses. On the other hand, underestimation patterns showed similar magnitudes for the same viral categories: 26.0% of DNA Eukaryota and 20.6% of RNA Eukaryota (non-influenza) assemblies were below -2 SD. In contrast, influenza genomes were almost never underestimated (<0.1%), confirming the model’s systematic overprediction for this group. When the model was asked to predict a “typical” Influenza virus (RNA, ss, eukaryotic host, mean GC% = 43.1%, mean genome size = 13.3 kb), it calculated an absolute Z-RNA sequence density value of 2.73 bps per kb, an overestimation of nearly 170-fold compared to the observed mean density of 0.016 bps per kb (relative model error > 17,000%).

Principal component analysis of dinucleotide enrichment profiles shows that influenza viruses occupy a restricted space relative to other viruses (**Fig. 5f**). The first two principal components explain most of the variance (PC1 = 38.8%, PC2 = 29%), with PC1 with higher AC, CG, and TA and lower CA, AG, and CC frequencies, while PC2 displays variation in GA and AT versus AA and CC. Most influenza genomes cluster close to the center, indicating they do not feature a strong bias towards specific dinucleotides. Coloring by Z-nucleic density suggests that higher density viruses cluster mostly in the PC1/PC2 positive space, where the enrichment of alternating purine-pyrimidine steps (AC, GC, AT, etc.) is higher (**Supp. Fig 1e**). Collectively, our analyses indicate that influenza viruses appear to have undergone strong evolutionary constraints that specifically purge Z-RNA-forming motifs, distinguishing them from all other viral groups.

## Discussion

In this first comprehensive study across 281,139 complete genomes of diverse lineages, we found that Z-nucleic acid sequence density is shaped by underlying sequence composition and selective pressures, and varies by orders of magnitude between organisms belonging to diverse taxonomic groups. Its strong correlation with GC content, especially in viral groups where GC content best predicts Z-DNA/Z-RNA density, underscores sequence composition as a major determinant. This is consistent with previous findings that CG and UA dinucleotides are strongly underrepresented in most RNA viral families (Di Giallonardo et al. 2017), which in our study exhibited the lowest Z-nucleic acid densities among all viral groups.

Enrichment analyses against dinucleotide-preserving controls demonstrate that Z-DNA tracts occur more frequently than expected in Bacteria, Archaea, and Eukaryota, and for specific taxa they seem to be enriched in functional regulatory elements. This highlights their role in key biological processes. Eukaryota is the most enriched group when examining Z-DNA prevalence around TSSs and TESs. *Homo sapiens* displayed high, but still lower enrichment around the TSS, than other primates, but seemed to feature higher Z-DNA density more upstream of the TSS. By contrast, Viruses displayed striking heterogeneity, with several RNA viral clades showing depletion, most notably observed in Influenza virus A and B genomes.

At a global scale, our findings reveal distinct evolutionary pressures shaping Z-nucleic acid formation in cellular versus viral genomes. In cellular life, Z-DNA appears to be an evolutionarily conserved mechanism for gene regulation and genomic plasticity, facilitating recombination and adaptive responses. On the other hand, several viral lineages feature a similar Z-nucleic acid presence, however specific groups of the Riboviria realm showed evidence of purifying selection against Z-forming motifs, perhaps to evade host ZBP1-mediated immune detection (Yin et al. 2025; Zhang et al. 2020). Similarly, SARS-CoV-2 and multiple RSV strains (both belonging to the same kingdom as Influenza viruses) exhibit very low or zero Z-RNA density. Conversely, viral species with a previously-described high Z-DNA potential like the Herpes Simplex Virus (Yin et al. 2025), showed a very high Z-DNA density (> 200 bps per kb). The intricate dynamics in Z-nucleic acid motifs further illustrate how structural avoidance of Z-forms can become fixed, providing a selective advantage in host-pathogen interactions. In other viral clades something similar may be achieved by different means. For example, some species in lineage-specific groups such as Nucleocytoviricota, Poxviruses, Cyprinid herpesviruses, and Asfarviridae, encode their own Zα-containing proteins that compete with host Z-DNA/-RNA sensors like ZBP1 to achieve immune evasion (Romero et al., 2024). Z-DNA structure and topology can indeed change rapidly, as these regions are more prone to radiation and oxidative damage than canonical B-DNA. They may be less susceptible to certain lesions like interstrand crosslinks, but their altered base positioning hinders repair efficiency (N7- or O6-methylguanine less efficiently repaired in Z-DNA) (Zhao et al. 2010). This makes Z-DNA both a source and a consequence of genomic instability; Z-DNA contributes to genomic instability by promoting DNA breaks and error-prone repair, while DNA damage, repair, and chromatin dynamics in turn influence the formation and topology of Z-DNA regions.

Although Z-RNA duplexes were not believed to form under natural conditions, they are now emerging as a new pathogen-associated molecular pattern. This along with Z-DNA’s implication in thymic T-cell tolerization (Fang et al. 2024) establishes Z-nucleic acids as relevant molecular structures in immunology. While most antiviral drug development has focused on G4 DNA, such data suggest that targeting Z-nucleic acids could similarly disrupt viral replication and enhance host defense, as observed with ZBP1 activation by natural compounds (Brázda et al. 2025). However, in addition to its role in viral clearance, ZBP1-mediated immune responses have been shown to amplify inflammation and tissue injury through activating multiple cell death associated pathways (Elsharkawy et al. 2025); thus, further studies in this area are warranted.

Altogether, these findings redefine the landscape of Z-DNA-forming potential across biological diversity. They reveal that while Z-DNA and Z-RNA motifs are widespread and often functionally enriched in cellular organisms, they are selectively suppressed in particular viral lineages. Future studies will be critical to elucidate the balance between the roles of Z-DNA in genome regulation, instability, and adaptation across the tree of life.

## Material and methods

### Data collection

Complete genomic assemblies were downloaded from the NCBI GenBank and RefSeq repositories on 09-02-2025 (Benson et al. 2013; O’Leary et al. 2016). Duplicate accessions were removed, choosing RefSeq over GenBank, as RefSeq entries are highly curated and quality-controlled by NIH. Each record was parsed to extract assembly descriptors including organism name, infraspecific name, taxonomy identifier, genome size and GC content. Genome size was manually corrected with a bash script that removed -N stretches longer than 10 nucleotides from the total base count. We also manually included additional telomere-to-telomere (T2T) assemblies beyond those downloaded by default from NCBI, drawing from various public sources (GenomeArk: https://github.com/genomeark/genomeark-update/, GigaScience Database: https://doi.org/10.5524/102657, Figshare repository: https://doi.org/10.6084/m9.figshare.25244542.v2, and T2T-Primates consortium (Yoo et al. 2025). The T2T genomes included, among others, Primates, Rodentia, Carnivora, Artiodactyla, and Perissodactyla (mammals); Poales, Rosales, Cucurbitales, Fabales, Solanales, Sapindales, Apiales, and Santalales (flowering plants); Saccharomycetales and Hypocreales (fungi); Enterobacterales and Mycoplasmoidales (Βacteria); and several representatives of teleost fishes such as Cypriniformes, Osmeriformes, Labriformes, Gymnotiformes. Metadata for our viral assemblies, including host and genome types, were downloaded from NCBI on 09-10-2025 from https://www.ncbi.nlm.nih.gov/labs/virus/vssi/#/virus. Our final dataset comprised 281,139 unique organismal genomes across all major taxonomic groups. The dataset was cross validated using NCBI’s Taxonomy database to confirm the consistency of organismal names and identifiers, before extracting full taxonomic lineages for each assembly.

### Prediction of Z-DNA sequences with ZSeeker

Z-DNA sequences were predicted with the ZSeeker software (version 1.8) (Wang et al. 2025) across all complete genomes. Predictions were executed with all scoring and penalty settings left at their default values (https://github.com/Georgakopoulos-Soares-lab/ZSeeker). Subsequently, the potential Z-DNA-forming sequences of each assembly were mapped to their chromosomal coordinates and intersected with overlapping genomic features.

### Estimation of Z-DNA density across genomes and relative to genomic sub-compartments

For each species, Z-DNA density was calculated by dividing the total length of all predicted Z-DNA sequences by the genome size in base pairs, then multiplying the result by 1,000 or 1,000,000 to obtain per kilobase or megabase density values, respectively. For each assembly, Z-DNA density within a given genomic feature (CDS, exon, gene) was calculated as the average fraction of each individual feature covered by Z-DNA, rather than the total proportion of all bases in that feature class. This ensured that each feature contributed equally to the estimate, irrespective of its length.

### Simulated genomes and Z-DNA enrichment

To estimate the expected amount and density of Z-DNA-forming sequences, we generated simulated control genomes using the fasta-shuffle-letters utility from the MEME Suite collection of tools (version 5.5.8) (Bailey et al. 2015). For each genome, we shuffled each chromosome’s sequence independently using the -kmer 2 option, which preserves the original dinucleotide composition while randomizing higher-order sequence context. Preserving dinucleotide frequencies is standard practice in sequence analysis (Gesell and Washietl 2008) and is particularly relevant here because Z-DNA formation is strongly influenced by dinucleotide patterns, especially alternating purine-pyrimidine motifs (Sahayasheela et al. 2025). We did not merge or reorder chromosomes, nor shuffle across chromosome boundaries, to retain the original genome architecture.

For each simulated genome, we predicted Z-DNA-forming sequences using the same Z-Seeker configuration and parameters as applied to the original NCBI reference genomes. We then calculated Z-DNA density per kilobase for both the simulated and observed genomes and assessed enrichment or depletion using the following formula:

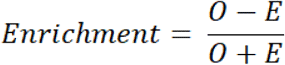

where *O* = observed Z-DNA density in the original genome, and *E* = expected Z-DNA density in the simulated genome. This metric ranges from −1 (complete depletion) to +1 (complete enrichment). In cases where both the original and simulated genomes contained no Z-DNA-forming sequences (*O* = *E* = 0), enrichment was defined as 0, indicating neither depletion nor enrichment. In cases where bound enrichment was not computed/needed, we calculated the fraction Obs./Exp. instead.

### Estimation of Z-DNA density relative to TSSs and TESs

For each assembly, we calculated the fraction of bases overlapping Z-DNA within a ±150 bp window around transcription start sites (TSSs) and transcription end sites (TESs). TSS and TES coordinates were defined as the first and last base pair of an annotated gene. Strand orientation was taken into account so that positions were aligned relative to the direction of transcription. For every gene, the proportion of Z-DNA coverage was computed at each position in the window, and these values were then averaged across all genes in the assembly to obtain genome-wide profiles of Z-DNA density relative to TSSs and TESs.

### Dinucleotide composition analysis

Dinucleotide frequencies were calculated using the EMBOSS version 6.6.0 suite program compseq (Rice et al. 2000). We applied:

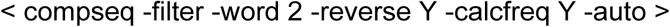

to each assembly, through a command-line interface. The “-word 2” option specifies calculation of dinucleotides, while “-reverse Y” counts occurrences in both strands. The “-calcfreq Y” option sets expected frequencies according to the observed mononucleotide composition of the genome, normalizing for differences in base composition (e.g. GC content). The program outputs observed counts, observed frequencies, expected frequencies, and observed-over-expected (O/E) ratios. The expected frequency of a dinucleotide XY is defined as:

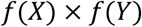

from the base composition. Therefore, the results represent whole-genome dinucleotide composition across all contigs/chromosomes within each assembly.

### Statistical and LASSO regression analyses

Statistical analyses used Mann–Whitney U tests (Bonferroni-adjusted) for unpaired and Wilcoxon signed-rank tests (FDR-adjusted) for paired comparisons with rank-biserial correlation coefficients reported as effect sizes. We applied LASSO (Least Absolute Shrinkage and Selection Operator) regression to model Z-DNA/Z-RNA motif density across viral genomes. The analysis was implemented in Python (scikit-learn, version 1.6.1) using a 5-fold cross-validated LassoCV estimator to automatically select the optimal regularization parameter (λ) minimizing prediction error. Predictors included G+C content, genome size (log₁₀-transformed), nucleic acid type (DNA/RNA), strand structure (ss/ds), and host type (Eukaryota/Prokaryota). Model performance was assessed by the mean cross-validated coefficient of determination (R²).

## Code availability

All relevant scripts and analysis code can be found on Github at this link: https://github.com/Georgakopoulos-Soares-lab/z-dna-taxonomic-analysis, while the ZSeeker outputs of all genome assemblies are archived in Zenodo at https://zenodo.org/records/17533316.

## Acknowledgements

Research reported in this publication was supported by the National Institute of General Medical Sciences of the National Institutes of Health under award number R35GM155468 and start-up funds awarded to I.G.S., and an R01, NIH/NCI (CA093729) to K.M.V..

## Supplementary figures

**Supplementary Figure 1:**
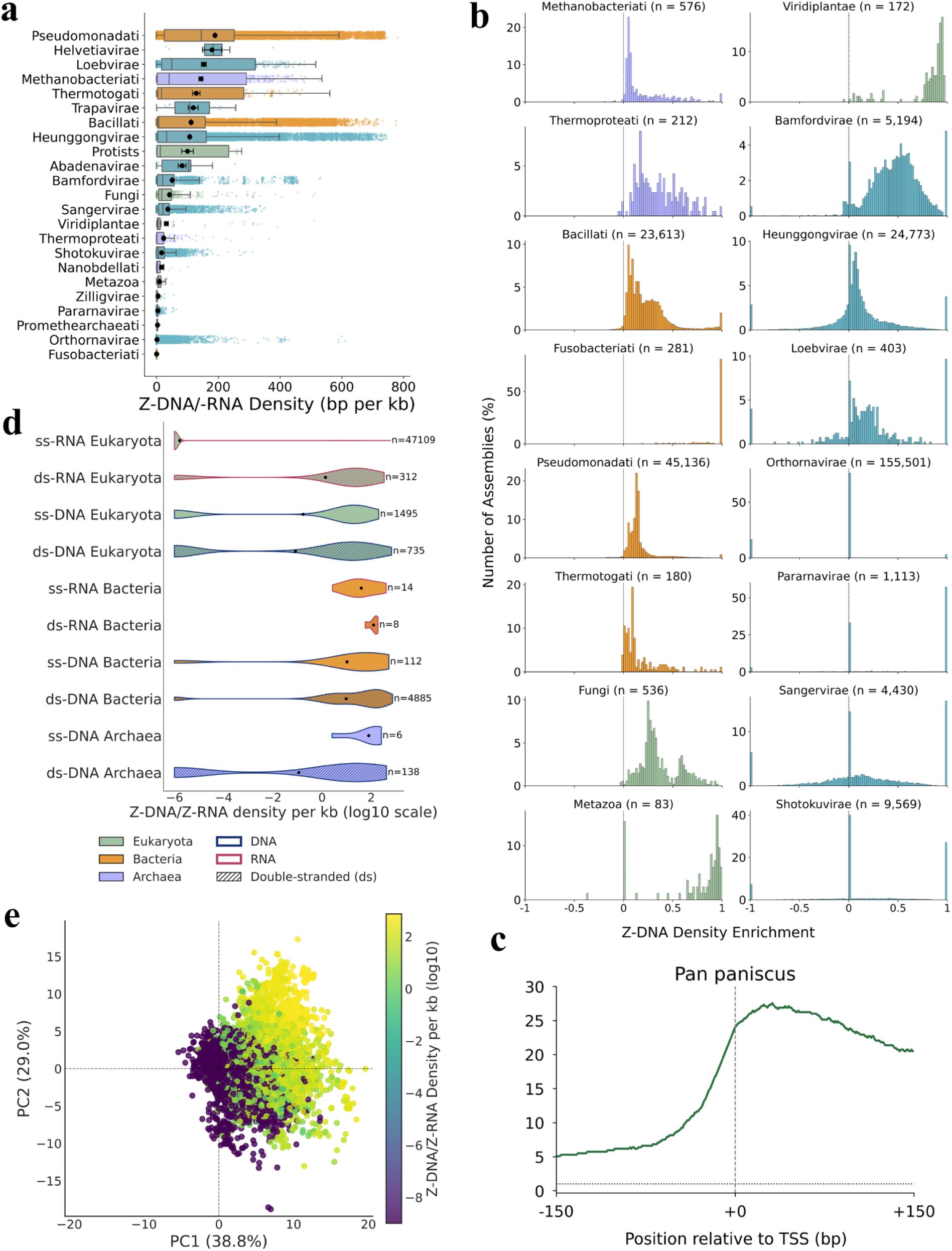
**a.** Raw Z-DNA density box plot of kingdoms, with mean and SEM error bars annotation. **b.** Bounded enrichment scores per kingdom. **c.** Log₂ fold-enrichment of Z-DNA around the TSS in *Pan paniscus*. **d.** Violin plot of Z-DNA/Z-RNA density across major viral groups, including strand, genome, and host type, averaged by unique taxonomy identifiers (n annotation). **e.** Principal component analysis in the dinucleotide enrichment space colored by log-transformed Z-DNA density.

## Supplementary Tables

**Supplementary Table 1:** Highest Z-DNA/-RNA genomes per superkingdom and across the dataset.

| Per superkingdom |  |  |
| --- | --- | --- |
| Superkingdom | Species (+ strain) | Density (bps per kb) |
| Archaea | *Halobaculum litoreum* DT92 | 535.9 |
| Bacteria | *uncultured beta proteobacterium* | 781.0 |
| Viruses | *Gordonia phage Flatwoods* | 744.3 |
| Eukaryota | *Malassezia furfur* | 512.2 |
| Globally |  |  |
| Bacteria | *uncultured beta proteobacterium* | 781.0 |
| Bacteria | *Cellulomonas iranensis* YLG372 | 775.0 |
| Bacteria | *Cellulomonas iranensis* ZJW-6 | 774.6 |
| Bacteria | *Gemmatirosa kalamazoonensis* KBS708 | 746.8 |
| Viruses | *Gordonia phage Flatwoods* | 744.3 |
| Viruses | *Gordonia Phage StorminNorm* | 744.2 |
| Viruses | *Gordonia phage Jeanie* | 742.7 |
| Viruses | *Gordonia phage McGonagall* | 742.6 |
| Bacteria | *Burkholderia pseudomallei* BPs116 | 739.4 |
| Viruses | *Gordonia phage Tangerine* | 739.3 |

**Supplementary Table 2:** Highest Z-DNA/-RNA predictions’ lengths and scores per superkingdom and across the dataset.

| Highest lengths per superkingdom |  |  |
| --- | --- | --- |
| Superkingdom | Species (+ strain) | Absolute length value |
| Archaea | *Cenarchaeum sp.* | 1,229 |
| Bacteria | *Cellvibrionales bacterium* | 26,493 |
| Eukaryota | *Neophocaena sunameri* | 65,589 |
| Viruses | *uncultured Caudovirales phage* | 2,008 |
| 5 highest lengths globally |  |  |
| Eukaryota | *Neophocaena sunameri* | 65,589 |
| Eukaryota | *Electrophorus electricus* | 65,316 |
| Eukaryota | *Arachis hypogaea* | 56,570 |
| Eukaryota | *Mus musculus C57BL/6* | 51,299 |
| Eukaryota | *Mus musculus C57BL/6J* | 50,172 |
| Highest scores per superkingdom |  |  |
| Superkingdom | Species (+ strain) | Absolute score value |
| Archaea | *uncultured Candidatus Cenarchaeum sp.* | 2,257.75 |
| Bacteria | *Mycobacterium marinum CCUG20998* | 23,162.75 |
| Eukaryota | *Electrophorus electricus* | 62,516.25 |
| Viruses | *uncultured Caudovirales phage* | 2,066.75 |
| 5 highest scores globally |  |  |
| Eukaryota | *Electrophorus electricus* | 62,516.25 |
| Eukaryota | *Arachis hypogaea* | 57,247.00 |
| Eukaryota | *Arachis hypogaea* | 48,182.00 |
| Eukaryota | *Equus asinus* | 36,632.50 |
| Eukaryota | *Pangasianodon hypophthalmus* | 36,437.25 |

## References

Azorin F, Nordheim A, Rich A. 1983. Formation of Z-DNA in negatively supercoiled plasmids is sensitive to small changes in salt concentration within the physiological range. EMBO J 2: 649–655.

Bacolla A, Tainer JA, Vasquez KM, Cooper DN. 2016. Translocation and deletion breakpoints in cancer genomes are associated with potential non-B DNA-forming sequences. Nucleic Acids Res 44: 5673–5688.

Bailey TL, Johnson J, Grant CE, Noble WS. 2015. The MEME Suite. Nucleic Acids Res 43: W39–49.

Balachandran S, Mocarski ES. 2021. Viral Z-RNA triggers ZBP1-dependent cell death. Curr Opin Virol 51: 134–140.

Beknazarov N, Konovalov D, Herbert A, Poptsova M. 2024. Z-DNA formation in promoters conserved between human and mouse are associated with increased transcription reinitiation rates. Sci Rep 14: 17786.

Benson DA, Cavanaugh M, Clark K, Karsch-Mizrachi I, Lipman DJ, Ostell J, Sayers EW. 2013. GenBank. Nucleic Acids Res 41: D36–42.

Brázda V, Bowater RP, Pečinka P, Bartas M. 2025. mGem: Noncanonical nucleic acid structures-powerful but neglected antiviral targets. mBio 16: e0273025.

Buzzo JR, Devaraj A, Gloag ES, Jurcisek JA, Robledo-Avila F, Kesler T, Wilbanks K, Mashburn-Warren L, Balu S, Wickham J, et al. 2021. Z-form extracellular DNA is a structural component of the bacterial biofilm matrix. Cell 184: 5740–5758.e17.

Cer RZ, Donohue DE, Mudunuri US, Temiz NA, Loss MA, Starner NJ, Halusa GN, Volfovsky N, Yi M, Luke BT, et al. 2013. Non-B DB v2.0: a database of predicted non-B DNA-forming motifs and its associated tools. Nucleic Acids Res 41: D94–D100.

Di Giallonardo F, Schlub TE, Shi M, Holmes EC. 2017. Dinucleotide Composition in Animal RNA Viruses Is Shaped More by Virus Family than by Host Species. J Virol 91. 10.1128/JVI.02381-16.

Elsharkawy A, Jahantigh HR, Guglani A, Stone S, Arora K, Kumar M. 2025. Virus-specific host responses and gene signatures following infection with major SARS-CoV-2 variants of concern: role of ZBP1 in viral clearance and lung inflammation. Front Immunol 16: 1557535.

Fang Y, Bansal K, Mostafavi S, Benoist C, Mathis D. 2024. AIRE relies on Z-DNA to flag gene targets for thymic T cell tolerization. Nature 628: 400–407.

Forni D, Pozzoli U, Cagliani R, Sironi M. 2024. Dinucleotide biases in the genomes of prokaryotic and eukaryotic dsDNA viruses and their hosts. Mol Ecol 33: e17287.

Freund AM, Bichara M, Fuchs RP. 1989. Z-DNA-forming sequences are spontaneous deletion hot spots. Proc Natl Acad Sci U S A 86: 7465–7469.

Georgakopoulos-Soares I, Morganella S, Jain N, Hemberg M, Nik-Zainal S. 2018. Noncanonical secondary structures arising from non-B DNA motifs are determinants of mutagenesis. Genome Res 28: 1264–1271.

Georgakopoulos-Soares I, Victorino J, Parada GE, Agarwal V, Zhao J, Wong HY, Umar MI, Elor O, Muhwezi A, An J-Y, et al. 2022. High-throughput characterization of the role of non-B DNA motifs on promoter function. Cell Genom 2. 10.1016/j.xgen.2022.100111.

Gesell T, Washietl S. 2008. Dinucleotide controlled null models for comparative RNA gene prediction. BMC Bioinformatics 9: 248.

Herbert A. (2019). Z-DNA and Z-RNA in human disease. Communications biology, 2, 7. 10.1038/s42003-018-0237-x

Ho PS, Ellison MJ, Quigley GJ, Rich A. 1986. A computer aided thermodynamic approach for predicting the formation of Z-DNA in naturally occurring sequences. EMBO J 5: 2737–2744.

Jovin TM, McIntosh LP, Arndt-Jovin DJ, Zarling DA, Robert-Nicoud M, van de Sande JH, Jorgenson KF, Eckstein F. 1983. Left-handed DNA: from synthetic polymers to chromosomes. J Biomol Struct Dyn 1: 21–57.

Kha DT, Wang G, Natrajan N, Harrison L, Vasquez KM. 2010. Pathways for double-strand break repair in genetically unstable Z-DNA-forming sequences. J Mol Biol 398: 471–480.

Kim YG, Lowenhaupt K, Maas S, Herbert A, Schwartz T, Rich A. 2000. The zab domain of the human RNA editing enzyme ADAR1 recognizes Z-DNA when surrounded by B-DNA. J Biol Chem 275: 26828–26833.

Kim Y-G, Muralinath M, Brandt T, Pearcy M, Hauns K, Lowenhaupt K, Jacobs BL, Rich A. 2003. A role for Z-DNA binding in vaccinia virus pathogenesis. Proc Natl Acad Sci U S A 100: 6974–6979.

Kouzine F, Wojtowicz D, Baranello L, Yamane A, Nelson S, Resch W, Kieffer-Kwon K-R, Benham CJ, Casellas R, Przytycka TM, et al. 2017. Permanganate/S1 Nuclease Footprinting Reveals Non-B DNA Structures with Regulatory Potential across a Mammalian Genome. Cell Syst 4: 344–356.e7.

Krall JB, Nichols PJ, Henen MA, Vicens Q, Vögeli B. 2023. Structure and Formation of Z-DNA and Z-RNA. Molecules 28. 10.3390/molecules28020843.

Lafer EM, Möller A, Nordheim A, Stollar BD, Rich A. 1981. Antibodies specific for left-handed Z-DNA. Proc Natl Acad Sci U S A 78: 3546–3550.

Li TT, D’Amico A, Christensen L, Vasquez KM. 2025. Effects of Aging on Z-DNA-Induced Genetic Instability In Vivo. Genes (Basel) 16. 10.3390/genes16080942.

McKinney JA, Wang G, Mukherjee A, Christensen L, Subramanian SHS, Zhao J, Vasquez KM. 2020. Distinct DNA repair pathways cause genomic instability at alternative DNA structures. Nat Commun 11: 236.

Nordheim A, Lafer EM, Peck LJ, Wang JC, Stollar BD, Rich A. 1982. Negatively supercoiled plasmids contain left-handed Z-DNA segments as detected by specific antibody binding. Cell 31: 309–318.

Nordheim A, Rich A. 1983. The sequence (dC-dA)n X (dG-dT)n forms left-handed Z-DNA in negatively supercoiled plasmids. Proc Natl Acad Sci U S A 80: 1821–1825.

Oh D-B, Kim Y-G, Rich A. 2002. Z-DNA-binding proteins can act as potent effectors of gene expression in vivo. Proc Natl Acad Sci U S A 99: 16666–16671.

O’Leary NA, Wright MW, Brister JR, Ciufo S, Haddad D, McVeigh R, Rajput B, Robbertse B, Smith-White B, Ako-Adjei D, et al. 2016. Reference sequence (RefSeq) database at NCBI: current status, taxonomic expansion, and functional annotation. Nucleic Acids Res 44: D733–45.

Peck LJ, Nordheim A, Rich A, Wang JC. 1982. Flipping of cloned d(pCpG)n.d(pCpG)n DNA sequences from right- to left-handed helical structure by salt, Co(III), or negative supercoiling. Proc Natl Acad Sci U S A 79: 4560–4564.

Peck LJ, Wang JC. 1983. Energetics of B-to-Z transition in DNA. Proc Natl Acad Sci U S A 80: 6206–6210.

Rice P, Longden I, Bleasby A. 2000. EMBOSS: the European Molecular Biology Open Software Suite. Trends Genet 16: 276–277.

Rich A, Zhang S. 2003. Timeline: Z-DNA: the long road to biological function. Nat Rev Genet 4: 566–572.

Rothan HA, Arora K, Natekar JP, Strate PG, Brinton MA, Kumar M. 2019. Z-DNA-Binding Protein 1 Is Critical for Controlling Virus Replication and Survival in West Nile Virus Encephalitis. Front Microbiol 10: 2089.

Run Y, Tavakoli M, Zhang Y, Vasquez KM, Zhang W. 2025. Formation and biological implications of Z-DNA. Trends Genet. 10.1016/j.tig.2025.07.006.

Sahayasheela VJ, Ooga M, Kumagai T, Sugiyama H. 2025. Z-DNA at the crossroads: untangling its role in genome dynamics. Trends Biochem Sci 50: 267–279.

Schroth GP, Chou PJ, Ho PS. 1992. Mapping Z-DNA in the human genome. Computer-aided mapping reveals a nonrandom distribution of potential Z-DNA-forming sequences in human genes. J Biol Chem 267: 11846–11855.

Schwartz T, Rould MA, Lowenhaupt K, Herbert A, Rich A. 1999. Crystal structure of the Zalpha domain of the human editing enzyme ADAR1 bound to left-handed Z-DNA. Science 284: 1841–1845.

Shin S-I, Ham S, Park J, Seo SH, Lim CH, Jeon H, Huh J, Roh T-Y. 2016. Z-DNA-forming sites identified by ChIP-Seq are associated with actively transcribed regions in the human genome. DNA Res 23: 477–486.

Subramani VK, Kim KK. 2023. Characterization of Z-DNA Using Circular Dichroism. Methods Mol Biol 2651: 33–51.

Umerenkov D, Herbert A, Konovalov D, Danilova A, Beknazarov N, Kokh V, Fedorov A, Poptsova M. 2023. Z-flipon variants reveal the many roles of Z-DNA and Z-RNA in health and disease. Life Sci Alliance 6. 10.26508/lsa.202301962.

Wang AH, Quigley GJ, Kolpak FJ, Crawford JL, van Boom JH, van der Marel G, Rich A. 1979. Molecular structure of a left-handed double helical DNA fragment at atomic resolution. Nature 282: 680–686.

Wang G, Christensen LA, Vasquez KM. 2006. Z-DNA-forming sequences generate large-scale deletions in mammalian cells. Proc Natl Acad Sci U S A 103: 2677–2682.

Wang G, Christensen L, Vasquez KM. 2023. Methods to Study Z-DNA-Induced Genetic Instability. Methods Mol Biol 2651: 227–240.

Wang G, Mouratidis I, Provatas K, Chantzi N, Patsakis M, Georgakopoulos-Soares I, Vasquez KM. 2025. ZSeeker: an optimized algorithm for Z-DNA detection in genomic sequences. Brief Bioinform 26. 10.1093/bib/bbaf240.

Wang G, Vasquez KM. 2023. Dynamic alternative DNA structures in biology and disease. Nat Rev Genet 24: 211–234.

Wang G, Vasquez KM. 2007. Z-DNA, an active element in the genome. Front Biosci 12: 4424–4438.

Wong B, Chen S, Kwon J-A, Rich A. 2007. Characterization of Z-DNA as a nucleosome-boundary element in yeast Saccharomyces cerevisiae. Proceedings of the National Academy of Sciences of the United States of America 104: 2229.

Xie KT, Wang G, Thompson AC, Wucherpfennig JI, Reimchen TE, MacColl ADC, Schluter D, Bell MA, Vasquez KM, Kingsley DM. 2019. DNA fragility in the parallel evolution of pelvic reduction in stickleback fish. *Science (New York*, NY) 363. https://pubmed.ncbi.nlm.nih.gov/30606845/ (Accessed July 5, 2025).

Yin C, Fedorov A, Guo H, Crawford JC, Rousseau C, Zhong X, Williams RM, Gautam A, Koehler HS, Whisnant AW, et al. 2025. Host cell Z-RNAs activate ZBP1 during virus infections. Nature. 10.1038/s41586-025-09705-5.

Yoo D, Rhie A, Hebbar P, Antonacci F, Logsdon GA, Solar SJ, Antipov D, Pickett BD, Safonova Y, Montinaro F, et al. 2025. Complete sequencing of ape genomes. Nature 641: 401–418.

Zhang T, Yin C, Boyd DF, Quarato G, Ingram JP, Shubina M, Ragan KB, Ishizuka T, Crawford JC, Tummers B, et al. 2020. Influenza Virus Z-RNAs Induce ZBP1-Mediated Necroptosis. Cell 180: 1115–1129.e13.

Zhang T, Yin C, Fedorov A, Qiao L, Bao H, Beknazarov N, Wang S, Gautam A, Williams RM, Crawford JC, et al. 2022. ADAR1 masks the cancer immunotherapeutic promise of ZBP1-driven necroptosis. Nature 606: 594–602.

Zhao J, Bacolla A, Wang G, Vasquez KM. 2010. Non-B DNA structure-induced genetic instability and evolution. Cell Mol Life Sci 67: 43–62.

